# The assessment of therapeutic autophagy inhibition in NF1 mutated tumor cells

**DOI:** 10.1101/2024.11.12.623325

**Authors:** Shadi Zahedi, Eliza Baird-Daniel, Kyle B. Williams, Rory L. Williams, Andrew Morin, Michelle Crespo, Emil Bratbak, Kendra Jones, Rajeev Vibhakar, Nicolas K. Foreman, David A. Largaespada, Jean Mulcahy Levy

## Abstract

MAPK pathway activation is found across central nervous system tumors. Neurofibromin 1 (NF1) protein is a GTPase-activating protein and tumor suppressor that negatively regulates RAS protein activity. A NF1 loss of function mutation results in MAPK pathway upregulation and is associated with pediatric gliomas. We previously demonstrated autophagy inhibition improves tumor response to BRAF inhibition. Here we investigate the role of autophagy in NF1 cells in the presence or absence of MEK inhibition (MEKi). NF1 wild type (WT) and NF1 knockout (KO) (HSC1λ, immortalized human Schwann) cells were evaluated. NF1 loss was confirmed, and cell growth was monitored by Incucyte. MAPK/ERK pathway upregulation and autophagy were evaluated via Western blot analysis and RAS pull-down. Effects of MEKi, autophagy inhibition (pharmacologic and genetic), and combination therapy were evaluated via short- and long-term growth assays. NF1 KO cells exhibited MAPK pathway upregulation and increased growth. Additionally, NF KO cells demonstrated increased autophagic activity. Loss of NF1 resulted in increased sensitivity to MEKi and autophagy inhibition alone and in combination. We also demonstrate that the autophagy inhibitor chloroquine (CQ) can improve MEKi sensitivity in a patient harboring a somatic mutation of NF1. This combination is currently being evaluated in clinical trial (NCT04201457).

## Introduction

Neurofibromatosis Type 1 (NF1) is an autosomal dominant disorder that arises due to a mutation in the NF1 gene encoding neurofibromin. Neurofibromin contains a central domain homologous to Ras-GTPase-activating proteins (Ras-GAPs) which function as negative regulators of Ras [1]. The prevalence of NF1 is estimated to be ∼1/3000 individuals [2, 3]. Germline NF1 is associated with plexiform neurofibromas - benign nerve sheath tumors that occur on peripheral and cranial nerves. Additional classic findings include café-au-lait macules, and Lisch nodules [4]. The condition is also associated with neoplasms in both the peripheral nervous system (PNS) and central nervous system (CNS). This includes plexiform neurofibromas which may cause disfigurement or compromise in peripheral nerve function, optic pathway gliomas, brainstem gliomas, and malignant peripheral nerve sheath tumors (MPNST)- the leading cause of early death in this patient population [5–8]. Tumors can also harbor somatic mutations of NF1 including high grade gliomas [9–11]. RAS hyperactivation due to NF1 loss results in downstream dysregulation of growth and survival pathways such as mitogen-activated protein kinase (MAPK) and mammalian target of rapamycin (mTOR) pathways [12]. Thus, utilizing kinase inhibitors to down regulate these pathways offers promising treatment opportunities. Imatinib mesylate has been shown to effectively decrease tumor volume in a phase II trial [13]. Additionally, mTOR inhibition via sirolimus has demonstrated increased time to progression in inoperable plexiform neurofibromas [14]. The MAPK pathway is a primary pathway of interest in NF1-mutated tumors. While dysregulation of downstream Raf/MEK/ERK pathway has been shown to induce negative feedback loops, ERK remains active, supporting how MEK plays a critical role in NF1 tumor growth. Pharmaceutical inhibition of MEK (MEKi) has demonstrated reduction in NF1 mutated tumor growth and decreased expression of downstream proliferation-associated genes [15]. Multiple clinical trials have also highlighted the benefit of MEKi in NF1 associated tumors [16–18].

Despite the development of these promising biologic therapies, there is well-documented emergence of acquired resistance to kinase inhibitor therapy in diverse cancer types. Tumors with described dysregulation of the MAPK pathway specifically have been found to develop multiple routes of acquired resistance to kinase inhibitor therapy, and resistance mechanisms are identified in a majority (58%) of tumors [19]. Additionally, tumors have often been found to develop more than one mechanism of resistance -- including both BRAF amplification and MEK mutation [20, 21]. Both primary and acquired resistance to MEKi via crosstalk with other pathways and feedback networks have also emerged as a barrier to chemotherapeutic efficacy [22, 23]. Thus, there is a great demand for methods to increase chemosensitivity in MAPK-dysregulated neoplasms such as those found in NF1.

Autophagy inhibition as a combination therapy is a potential strategy to overcome kinase inhibitor resistance. Autophagy is a heavily regulated catabolic process by which cellular material is transported to lysosomes for degradation and subsequent re-utilization for cellular processes [24, 25]. Autophagic activity is upregulated in stressful cellular conditions such as hypoxia, DNA damage, and nutrient deprivation states to increase availability of macromolecular metabolic precursors for cell survival. Moreover, core autophagy gene expression is dramatically upregulated in diverse cancer types suggesting that tumor cells become “autophagy-addicted” to overcome nutritional stresses associated with accelerated cellular proliferation and the relatively inhospitable conditions found in the tumor core [26].

Previous studies in adult gliomas with BRAF WT and PTEN mutations resistant to phosphatidylinositol 3-kinase to AKT to mammalian target of rapamycin (PI3K-AKT-mTOR) pathway inhibitors have demonstrated that inhibiting autophagy resulted in improved response to dual PI3K-mTOR inhibitors [27]. In melanoma tumor biopsies, inhibiting the upregulated autophagy induced through endoplasmic reticulum (ER) stress following a BRAF inhibitor (BRAFi) treatment with vemurafenib overcame drug resistance [28]. Increased autophagic activity has also been specifically described in MAPK dysregulated neoplasms [28, 29] as tumors with hyperactivation of ERK demonstrate increased levels of autophagy [30].

While autophagy is found to be constitutively upregulated in many tumors, it is observed to be further induced after initiation of chemotherapeutic agents. This suggests that autophagy deregulation not only allows for cellular proliferation in aggressive tumors, but may play a crucial role in chemo-evasion [25]. Intensified utilization of autophagy for cell survival and growth has been shown to be a critical aspect of acquired chemo-resistance in tumors with MAPK dysregulation [28, 29]. It has been shown that melanoma with higher autophagic index is less likely to respond to chemotherapeutic treatment [31].

Autophagy inhibition in MAPK dysregulated tumors has been demonstrated to increase chemosensitivity and improve clinical outcomes [29]. We have shown that, in pediatric CNS tumors containing a BRAF^V600E^ mutation, autophagy inhibition is effective in overcoming several mechanisms of chemoresistance [32]. The most utilized autophagy inhibitors to date are clinically approved antimalarial drug, chloroquine (CQ) and its derivatives such as hydroxy chloroquine (HCQ). CQ and HCQ are thought to block autophagy by accumulating inside endosomes leading to deacidification which in turn leads to impaired lysosomal enzymatic function [33]. Our group and others have shown the efficacy of using CQ for combating tumors that rely on autophagy for proliferation and survival [29, 34, 35] [36]. CNS tumors harboring BRAF^V600E^ mutation in the MAPK pathway have greater activation of autophagic pathways and demonstrate upregulation of autophagy when under stressors such as chemotherapy. Inhibition of autophagy in this subset of CNS tumors is associated with a greater decrease in tumor cellular viability than non-MAPK dysregulated controls [29]. Autophagy inhibition has yet to be explored in enhancing chemosensitivity to MEKi in NF1 mutated tumors.

Here, we show that high levels of autophagy are induced in NF1 KO cells. This results in increased dependence on autophagy and, thus, increased sensitivity to autophagy inhibition. Additionally, autophagy inhibition appears synergistic when combined with inhibition of MEK. We also demonstrate that the autophagy inhibitor CQ can improve the clinical efficacy of the MEK inhibitor Trametinib in a patient with a somatic mutation of NF1.

## Materials and Methods

### Study design

Experiments were designed to evaluate the hypothesis that CNS tumors with an upregulated MAPK pathway caused by alterations other than BRAF^V600E^, such as a NF1 mutation, will be equally autophagy dependent and this will result in synergy between MEKi (standard therapy) and autophagy inhibition, improving treatment for such tumors. To evaluate the effectiveness of therapies on cell growth and survival in both NF1 wild type and NF1 null cell lines, both long- and short-term growth and viability assays were utilized. We assessed dysregulated MAPK pathway in HSC1 lambda cells by western blot analyses and co-immunoprecipitation. The specificity to the autophagy pathway was evaluated with genetic inhibition studies. The final endpoints were defined prior to the start of each experiment. Details on replicates and statistical analysis are indicated in the figure legends.

### Cell culture and reagents

Primary Human Schwann cells were purchased from ScienCell (#1700, Carlsbad, CA). The cells were cultured in Schwann Cell Medium (SCM) (#1701). SCM consists of 500 ml of basal medium, 25 ml of FBS (#0025), 5 ml of Schwann Cell Growth Supplement (#1752) and 5 ml of penicillin/streptomycin solution (#0503). Isogenic Human Schwann cells (HSC1) with and without NF1 loss were kindly given to us by Dr. Largaespada [37–39]. The cells were cultured in DMEM (#10-013-CV) supplemented with 10% fetal bovine serum (FBS) (#S11150, Atlanta Biologicals, Flowery Branch, GA) containing 1% Penicillin/ streptomycin (#15070063, Thermo Fisher Scientific, Waltham, MA).

### Statistics

Statistical comparisons were completed by using an ordinary two-way (GraphPad Prism 7.04, RRID: SCR_002798) as indicated in the figure legends. A P-value of < 0.05 was considered statistically significant. The data shown are mean ± standard error of the mean (SEM) except where indicated.

### In vitro viability assays

For short-term viability assays, cells were seeded in 96-well plates at a density of 1000 cells/well followed by the treatment as indicated and incubated at 37 °C in 5% CO2. After 5 days of treatment, cell viability was determined by using CellTiter-Glo® Luminescent Cell Viability Assay according to the manufacturer’s protocol (#G7572, Promega Corporation, Madison, WI).

Long-term survival was assessed by using a colony formation assay. Cells were plated into 12-well plates at a density of 750 cells/well in accordance with the optimal growth condition of the cell line and incubated overnight at 37 °C in 5% CO2. Cells were treated with a IC50 dose of drugs as indicated and incubated for 10–14 days. Colonies were monitored and provided with fresh media with or without drugs every 3 days. When the control wells reached 80–85% confluence, the cells were fixed, stained with 0.4% crystal violet, and quantified by solubilization in 33% (v/v) acetic acid with A540 absorbance assessed.

### Incucyte growth measurement assay

Cells were seeded in 96-well plates (Costar, Corning, NY) at a density of 1000 cells/well. Cells were cultured at 37° and 5% CO2 and monitored using an Incucyte Zoom (BioScience Inc., Ann Arbor, MI). Cells were then treated as described and monitored. Images were captured at 4 hr intervals from four separate regions per well using a 10x objective. Each experiment was done in triplicate and growth curves were created from percent confluence measurements based on cell count per well.

### Western blot analysis and antibodies

Cell lysates were harvested after treatments and time points indicated by using RIPA buffer (Sigma, St. Louis, MO) with protease inhibitor cocktails (Roche, Indianapolis, IN). Samples were boiled for 10 min at 95 °C, and they were resolved by SDS-PAGE. Membranes were blocked with 5% dry nonfat milk in TBS-Tween for 1 h at room temperature and probed with primary antibodies at the manufacturer’s recommended concentrations. The primary antibodies used were NF1 (#ab17963); Phospho-p44/42 MAP kinase (pERK1/2) (#9101S) (Cell Signaling, Danvers, MA); p44/42 MAP kinase (Erk1/2) (#9102) (Cell Signaling, Danvers, MA); LC3 (#NB100–2220) (Novus Biologicals, Littleton, CO); RAS (#8832); ATG5 (#12994S) (Cell Signaling, Danvers, MA). Anti-β-actin (#12262, RRID: AB_2566811) (Cell Signaling, Danvers, MA) was used as the protein loading control. The secondary antibody used was anti-rabbit IgG (#7074S) (Cell Signaling, Danvers, MA). The results of western blots were assessed by comparing the intensity of bands by using Image J.

### RAS pull down assay

Cell lysates (500 µl at 1 mg/ml) were treated with GTPγS (positive control) or GDP (negative control) to activate or inactivate Ras. GST-Raf1-RBD fusion protein was used to bind the activated form of GTP-bound Ras. GTP-bound Ras was then immunoprecipitated with glutathione resin. Western blot analysis of cell lysate and eluted samples was performed using a RAS mouse mAb. Anti-mouse IgG, HRP-linked Antibody #7076 was used as the secondary antibody.

### ShRNA transfection

A pLKO.1-puro lentiviral vector from the RNAi Consortium (TRC; Sigma-Aldrich) was utilized with smallhairpin RNAs (shRNAs). TRC numbers for shRNAs used were as follows: ATG5 (#474), the nontarget (#SHC016). Cells were transduced with a lentivirus by using 8 µg/ml polybrene and harvested 3 days following transduction. The level of target gene knockdown and its effect on autophagic flux was determined via western blotting.

### MRI images

MRI images were obtained using standard protocols on a Siemens 1.5 T Avanto scanner.

## Results

### NF1 KO results in increased growth in HSC1 lambda cells

To assess the impact of NF1 loss on cell growth, cells were plated and monitored by using IncucyteZoom. Real-time measurements of cell growth over time (Fig. 1A-D) demonstrated a significant increase in the growth rate in NF1 KO cells compared to NF1 WT cells (Fig. 1E). On day four of monitoring, NF1 null cells exhibited 80.69% confluency compared to NF1 WT cells, which only showed a 55.18% confluency (Fig. 1C, D). To confirm a difference in NF1 protein expression between the cell lines, we performed a western blot analysis (Fig. 1F, Supplementary Figure 1) that confirmed loss of NF1 protein in NF1 KO cells.

**Figure 1.**
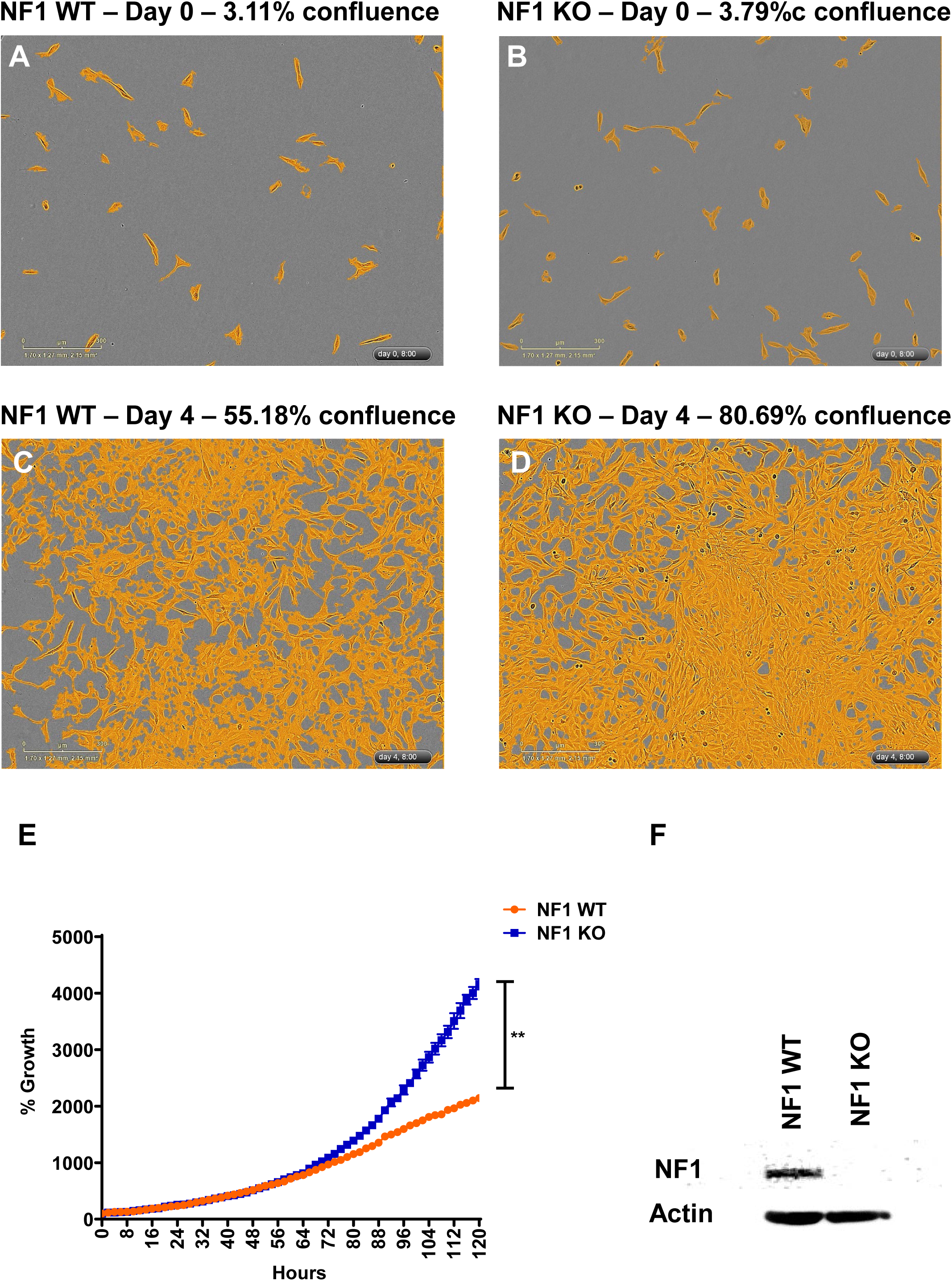
NFI KO cells demonstrate increased growth compared NF1 WT cells. **A-D)** Phase contrast images of NF1 WT and NF1 KO cells over a period of 120 hours shown with and without confluence mask (orange). **E)** Percent growth overtime in NF1 WT and NF1 KO cells. Growth measured by continuous incucyte monitoring (Two-way ANOVA; mean ± s.e.m., *n* = 2. \**p* < 0.05). **F)** Representative western showing the effectiveness of NF1 RNAi. Original blot is presented in Supplementary Figure 1.

### NF1 null cells showed MAPK pathway dysregulation and increased sensitivity to MEK inhibition

To test the effect of NF1 KO on regulating the MAPK pathway, we aimed to assess both the upstream and downstream NF1 targets within the pathway. NF1 is known to downregulate RAS. Hence, we measured activated RAS protein levels in NF1 WT and NF1 KO via a pull-down assay. The western blot analysis showed higher levels of activated RAS (GTP-bound) in the NF1 KO cells compared to NF1 WT as expected (Fig. 2A, Supplementary Figure 2A). To assess the downstream effects of NF1 loss, we measured ERK phosphorylation in NF1 WT and NF1 KO cells. Western blot analysis revealed an increase in phosphorylated ERK levels in NF1 KO cells compared to NF1 WT cells (Fig. 2B, C, Supplementary Figure 2B). These results suggest MAPK pathway hyperactivation in the NF1 KO cells compared to NF1 WT cells.

**Figure 2.**
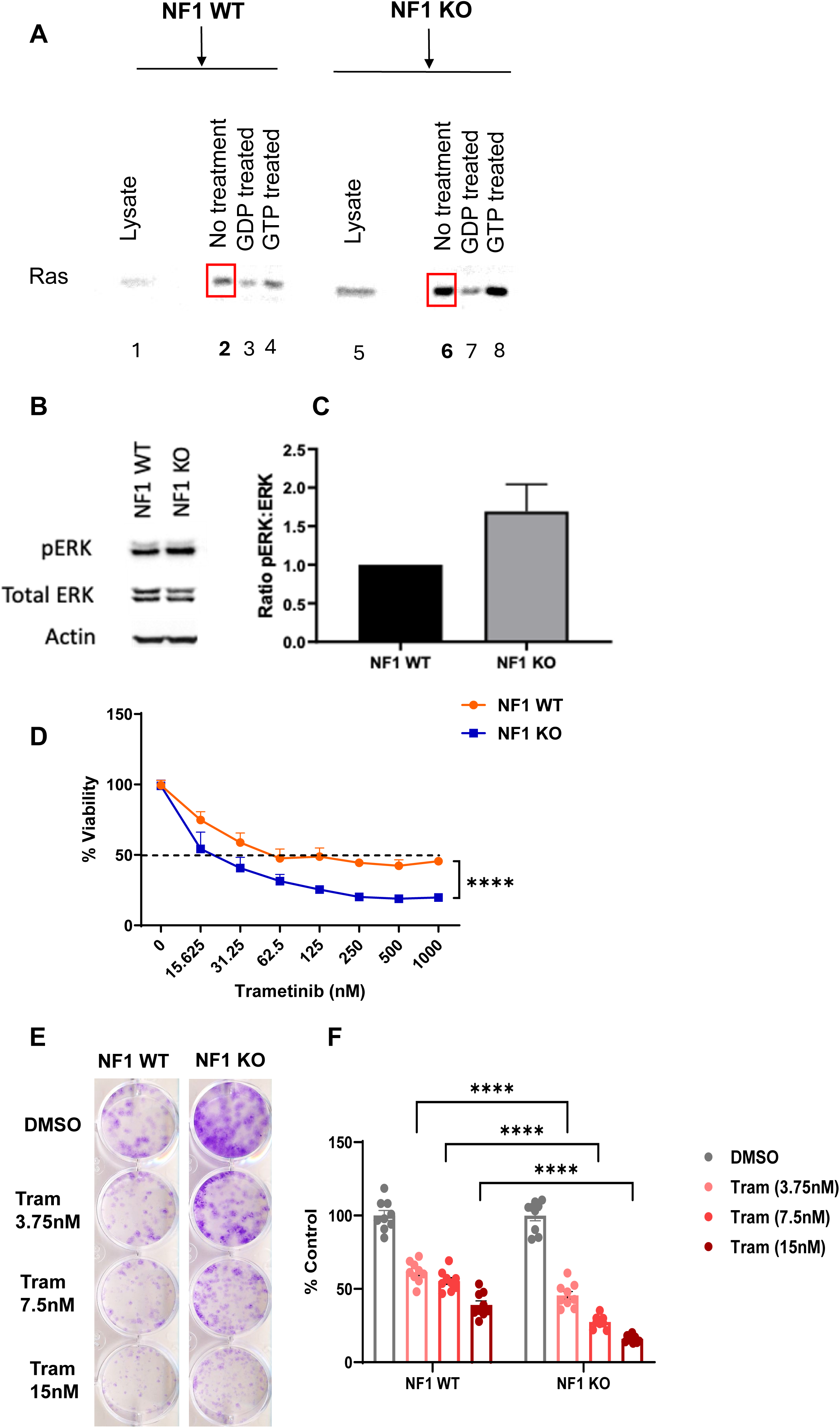
NF1 KO cells demonstrate MAPK pathway dysregulation and increased sensitivity to MEKi. **A)** Western blot depicting higher levels of activated (GTP-bound) Ras in the NF1 KO compared to NF1 WT (left). **B)** A representative western blot depicting phosphorylation of ERK in the NF1 KO compared to the NF1 WT. Original blot is presented in Supplementary Figure 2B. **C)** Bargraph showing the calculated ratio of phosphorylated-ERK:ERK with the NF1 KO cells showing greater phosphorylation. **D)** Effect of MEKi on short-term viability in NF1 WT and NF1 KO cells treated with increasing doses of Trametinib for 5 days. Viability was determined by using the CellTiter Glo assay (Two-way ANOVA; mean ± s.e.m., n = 3. *p < 0.05) **E, F**) Representative long-tern clonogenic assays (E) and quantified collated data (F) of cells treated with Trametinib as indicated (Two-way ANOVA; mean ± s.e.m., n = 3. *p < 0.05).

Next we tested whether our NF1 KO cells showed greater sensitivity to MEK inhibition (MEKi) compared to WT cells, as previously shown in NF1 mutated tumors [15]. This was examined through both short- and long-term viability and growth assays. For short-term viability analysis, cells were treated with increasing doses trametinib, a MEK inhibitor that is FDA approved to treat pediatric low-grade glioma. Endpoint cell viability measurements demonstrated a significant dose-dependent decrease in cell viability in NF1 KO cells compared to NF1 WT cells (Fig. 2D). In addition, clonogenic long-term growth assays indicated a significant dose-dependent sensitivity in NF1 KO compared to NF1 WT cells (Fig. 2E, F). Our data suggests an upregulated MAPK pathway in NF1 KO cells, which appears to enhance cell sensitivity toward MEK inhibition.

### Increased autophagic activity is seen in NF1 KO cells at baseline and in response to stress

To evaluate the role of autophagy in NF1 null cells, we assessed autophagic activity at baseline (Fig. 3A, B, Supplementary Figures 3A, B) and under stress conditions (Fig. 3C, D, Supplementary Figure 3C, D) in NF1 WT and NF1 KO cells. Representative western blots demonstrate a greater accumulation of LC3 II at lower CQ concentrations in NF1 KO cells compared to NF1 WT cells (Fig. 3A, Supplementary Figure 3A). Furthermore, we observed an earlier accumulation of LC3 II in NF1 KO cells following treatment with CQ (Fig. 3B, Supplementary Figure 3B). These data indicate higher autophagic activity in NF1 KO cells compared to NF1 WT cells. Under trametinib treatment (a stress condition) NF1 KO cells induced autophagy to a greater degree than WT cells (Fig. 3C, Supplementary Figure 3C). This was evaluated by measuring autophagic flux, comparing LC3II accumulation to the control. Western blot analyses revealed a significantly higher increase in LC3II accumulation upon autophagy inhibition while under trametinib treatment in NF1 KO cells (Fig. 3C, D, Supplementary Figure 3C, D). Overall, these data suggests that a lack of NF1 increases tumor cells’ dependence on autophagy, making them potentially more vulnerable to autophagy inhibition.

**Figure 3.**
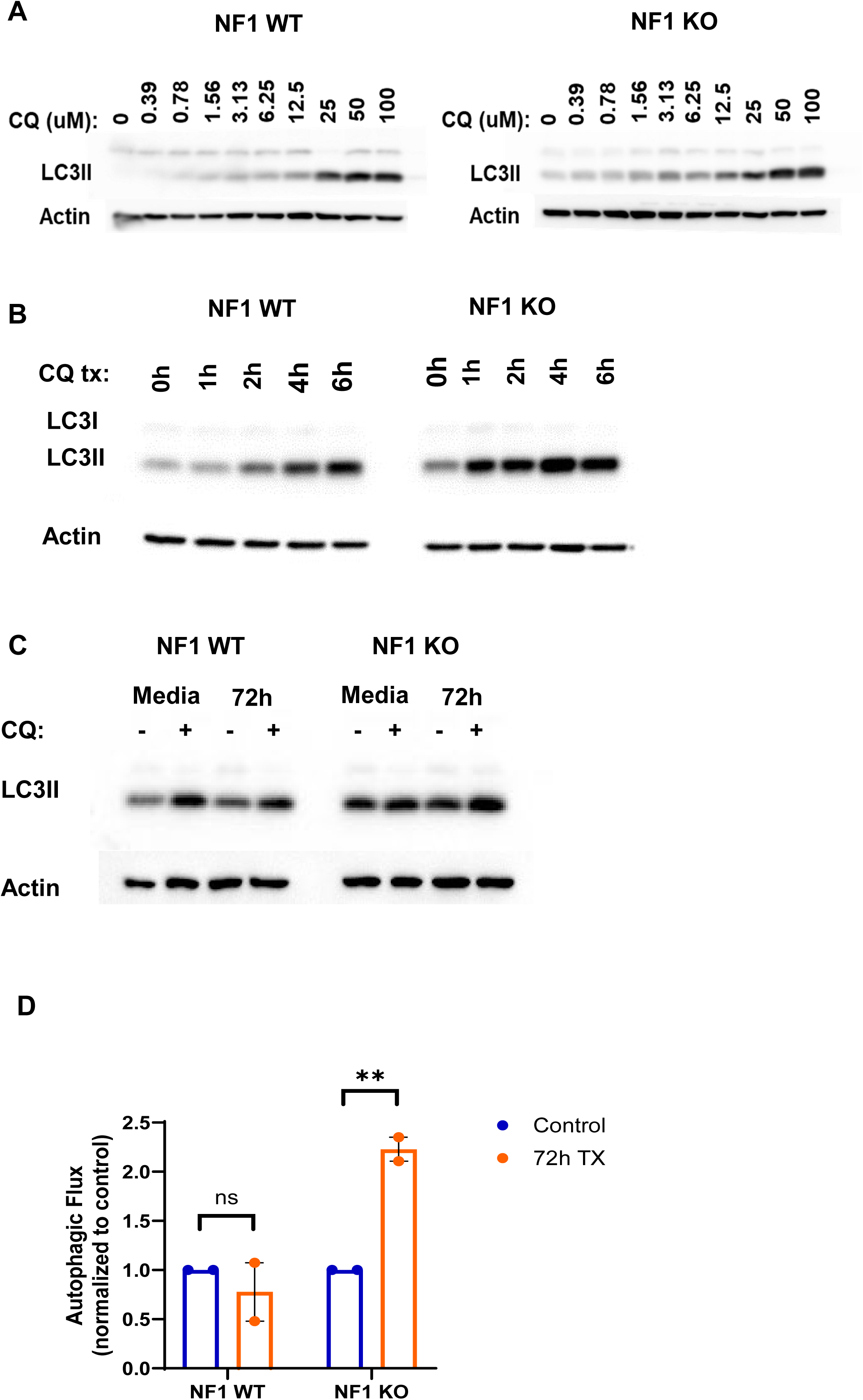
NF1 KO demonstrates increased autophagic activity at baseline and in response to stress. **A)** Representative western blot demonstrating a greater accumulation of LC3 in NF1 KO line at lower CQ concentrations. **B)** Representative western blot demonstrating earlier accumulation of LC3 in NF1 KO cells after treatment with 50uM CQ for 0, 1, 2, 4, and 6 hours. **C, D)** Representative western blot **(C)** and the quantification **(D)** demonstrating increased LC3 accumulation and autophagic flux in NF1 KO as compared to NF1 WT in response to Trametinib treatment with or without exposure to 50uM CQ for 4hrs (Unpaired t test; mean ± s.e.m., n = 2. *p < 0.05). Representative western showing the effectiveness of NF1 RNAi. Original blots are presented in Supplementary Figures 3A, B, C.

### NF1 KO cells exhibit increased sensitivity to autophagy inhibition both alone and in combination with MEK inhibition

We have previously shown pharmacologic inhibition of autophagy with CQ improved the effectiveness of BRAFV600E inhibition in BRAF mutated cells [29]. To assess whether autophagy inhibition could also support better the effectiveness of MEKi in NF1 mutated cells, we first examined short- and long-term viability and growth in both NF1 WT and NF1 KO cells following treatment with CQ. We initially compared NF1 KO cells to primary schwann cells to demonstrate a therapeutic window for CQ in NF1 KO cells compared to the parental cell. Results demonstrated NF1 KO cells were significantly more sensitive to increasing doses of CQ (Fig. 4A). Long-term clonogenic growth assays comparing the NF1 WT and NF1 KO cells again demonstrated NF1 KO cells grew faster as shown by DMSO control wells. Despite this, CQ was able to inhibit a larger number and percentage of cells in NF1 KO cells compared to NF1 WT cells (Fig. 4B, C). These data suggest NF1 KO cells are more vulnerable to autophagy inhibition. Additionally, we assessed the effect of autophagy inhibition in the presence or absence of MEKi. Although both WT and KO cells exhibited sensitivity to monotherapies, NF1 KO cells showed a significant decrease in cell viability when following MEKi and CQ combination therapy (Fig. 4D). Subsequent evaluation using a Bliss synergy model confirmed this synergistic effect (synergy maps highlight synergistic and antagonistic dose regions in red and green colors, respectively) (Fig. 4E).

**Figure 4.**
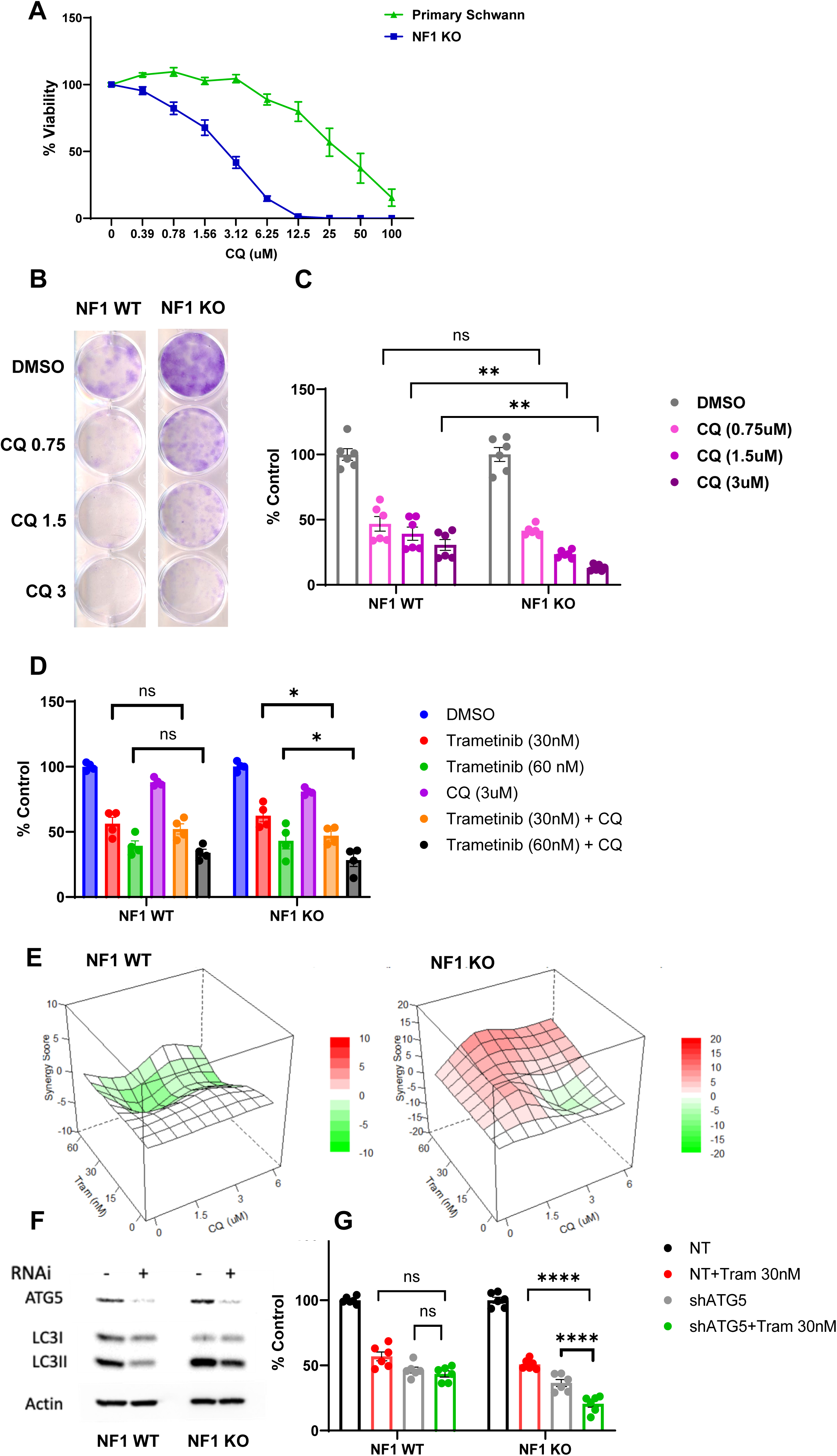
NF1 KO cells exhibit increased sensitivity to CQ both alone and in combination with Trametinib. **A)** Effect of CQ on short-term viability in primary Schwann and NF1 KO cells treated with increasing doses of CQ for 5 days. Viability was determined by using the CellTiter Glo assay. **B, C)** Representative long-tern clonogenic assays **(B)** and quantified collated data **(C)** of cells treated with CQ as indicated (Two-way ANOVA; mean ± s.e.m., n = 3. *p < 0.05). **D)** Percentage of cell viability as measured by CellTiter Glo (compared with control DMSO) following a 5-day exposure to trametinib and CQ alone and in combination (Two-way ANOVA; mean ± s.e.m., n = 3. *p < 0.05). **E)** Heatmap representation of the Bliss interaction index across the three-point dose range of Tram and CQ in NF1 WT and NF1 KO cells. Mean values of double biological experiments are shown. **F)** Representative westerns showing effectiveness of ATG5 RNAi and resultant decrease of LC3II. G) Percent viable cells, by Cell Titer-Glo (compared to control NT) following 120 hr of Trametinib drug therapy with and without RNAi of essential autophagy protein ATG5.

To ensure the benefits of CQ were due to autophagy inhibition and not secondary to off-target effects we knocked down ATG5 (Fig. 4F), an essential regulator of canonical autophagy, with and without MEKi. Confirmation that ATG5 knockdown inhibited autophagy was shown by a decrease in the autophagy marker LC3II (Fig. 4F). When ATG5 knockdown was present in NF1 WT cells, there was no significant change in response to MEKi with trametinib. In contrast, the addition of ATG5 knockdown in NF1 KO cells demonstrated a significant decrease in cell survival in combination treatment compared to either MEKi or autophagy inhibition alone (Fig. 4G). These data support that re-sensitization to MEKi seen in NF1 KO cells was due to autophagy inhibition and not another effect of CQ.

#### Autophagy inhibition decreases growth of brain tumor in a patient with a somatic NF1 mutation

An 11-year-old girl presented with several weeks of vomiting, somnolence, disconjugate gaze, dysarthria, and imbalance and was found to have a localized tumor of the left thalamus. She underwent debulking of her tumor at diagnosis, which showed a diffuse midline glioma, H3K27M-mutant. She was treated with focal radiation therapy, with which she had decreased surrounding T2 signal but increased central enhancement (Fig. 5A) compared to post-operative MRI. She then began maintenance temozolomide and lomustine. After two cycles, her MRI showed definitive tumor growth (Fig. 5B), so these were discontinued, and she was started on trametinib/chloroquine due to a somatic NF1 mutation identified in her tumor. After initial growth of the tumor six weeks later (Fig. 5C), she experienced a strong partial response to therapy by imaging (Fig. 5D, E) and has had no further progression in this primary tumor region over the subsequent three years.

**Figure 5.**
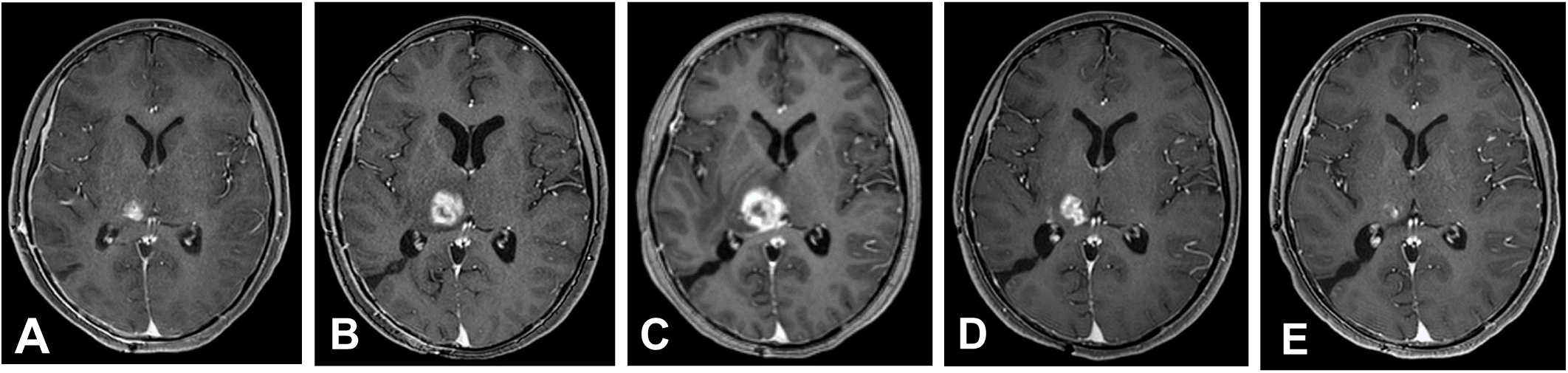
Axial T1-weighted post contrast images from sequential MRIs demonstrate progressively increased size of the enhancing left thalamic tumor on two-month interval scans after treatment with CCNU/TMZ from **(A)** to **(B)**. The patient was switched to trametinib/chloroquine starting after the progression seen in **(B).** After additional growth six weeks later seen in **(C)**, treatment response is depicted with decreased size of the enhancing mass on three-month interval scans **(D)** and **(E)**.

## Discussion

Previous studies have demonstrated that autophagy plays an important role in chemo-evasion in several tumor types with certain genetic mutations that result in upregulation of the MAPK pathway. In melanoma cell lines, autophagy inhibition overcame resistance due to the upregulation of ER stress-induced autophagy to BRAFi [24]. In adult GBM, the EGFRvIII mutation was identified as a marker of autophagy dependence. Treating EGFRvIII^+^ tumor cells with the autophagy inhibitor CQ reduced proliferation and compromised survival during stress conditions [35]. Additionally, our group demonstrated a reversal of clinical and radiographic disease progression after inhibiting autophagy in a BRAF^V600E^ ganglioglioma patient who had progressed on vemurafenib [25] and in a high-grade glioma patient with BRAF^V600E^ [34].

With the positive findings with autophagy inhibition in BRAF^V600E^ brain tumors, we questioned if other mechanisms of MAPK pathway activation would lead other brain tumor subgroups to have a similar response. Evaluation of non-BRAF V600E MAPK-activated tumors, such as tumors with NF1 mutation, would provide additional biomarkers of patients anticipated to have a response to autophagy inhibition. This has been shown as a possibility in pancreatic tumors with alternate mechanisms of activating the pathway [35, 36, 40, 41]. Our data suggests that brain tumors with loss of NF1 and secondary MAPK pathway activation are, like BRAF^V600E^ brain tumors [29, 34], also autophagy addicted and candidates for the addition of autophagy inhibition to current targeted therapies. This creates an expanded pool of patients that may benefit from clinical autophagy inhibition.

In the current study, we aimed to investigate the impact of autophagy inhibition on the NF1-mutated cell response alone and in combination with MEKi. First, we demonstrated that cells that lacked the NF1 gene showed an upregulation of the MAPK pathway with increased cellular growth and proliferation. When we assessed autophagic activity, we observed an increased basal autophagy in NF1 KO cells compared to WT cells. This agrees with our previous studies, where we demonstrated that cells with MAPK dysregulation due to BRAF^V600E^ have increased levels of autophagy and are, therefore, sensitive to autophagy inhibition. Our data also demonstrated a significant difference in growth and viability between WT cells and those lacking NF1 when treated with MEKi with and without autophagy inhibition. MEKi has proven effective in treating NF1-associated tumors [18]; however, similar to many targeted therapies, the development of resistance appears inevitable. And although MEKi has already been approved for treating NF1 and NF1-associated tumors, there has been no evidence of complete disease control with monotherapy [35].

Our combination studies demonstrated reduced viability in response to MEKi and both pharmacologic and genetic autophagy inhibition combination therapy in NF1 KO cells. This suggests autophagy inhibition may be the additional therapy needed to improve disease control in NF1 mutated tumors. Additional studies evaluating potential new therapies for these cells are also under investigation [39]. Despite the advantage of being highly controlled and focused, *in vitro* studies carry the disadvantage of uncertainty in depicting the effects that are seen in complex living organisms. Luckily, use of this approach has already been clinically successful in our group with a patient with diffuse midline glioma with a somatic mutation of NF1 where the combination of autophagy and MEK inhibition resulted in a strong, continued partial clinical response. Additional clinical evaluation of this combination is currently ongoing through the Pediatric Brain Tumor Consortium (NCT04201457).

## Data availability

All data analyzed in this study are available from the corresponding author on request.

## Author contributions

SZ and JML designed the project; JML directed all work related to the project and data analysis; KW, LW, DAL provided Isogenic Human Schwann cells (HSC1) and participated in data and manuscript review; SZ, EBD, MC conducted experiments; SZ analyzed data and prepared figures; SZ and EBD wrote the initial manuscript; JML, EBD, MC, AM, EB, KJ, NF, and RV participated in data and manuscript review and edit.

