## Supplementary Figures for "The assessment of therapeutic autophagy inhibition in NF1 mutated tumor cells"

Supplementary Figure 1.

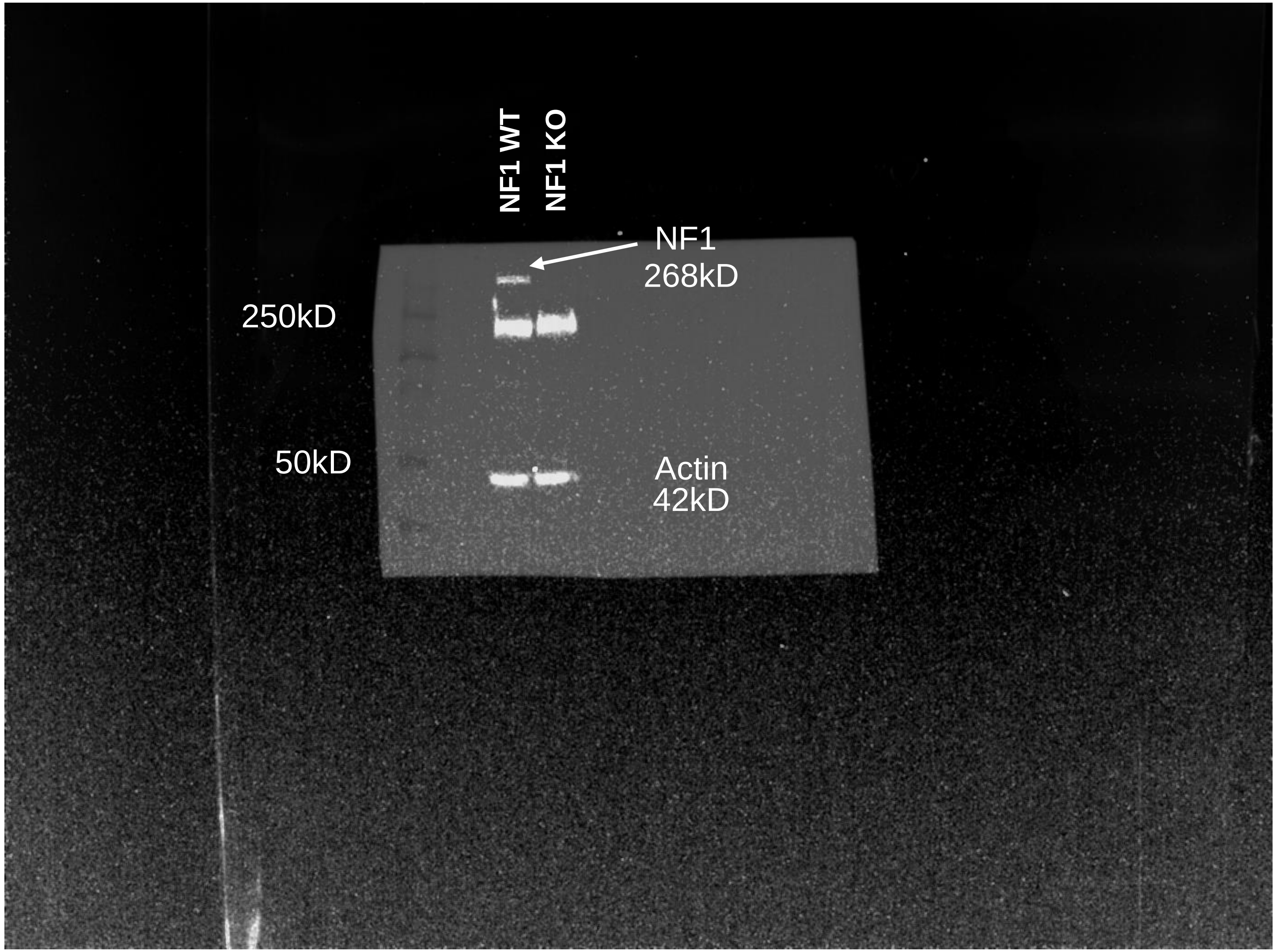

Supplementary Figure 2A.

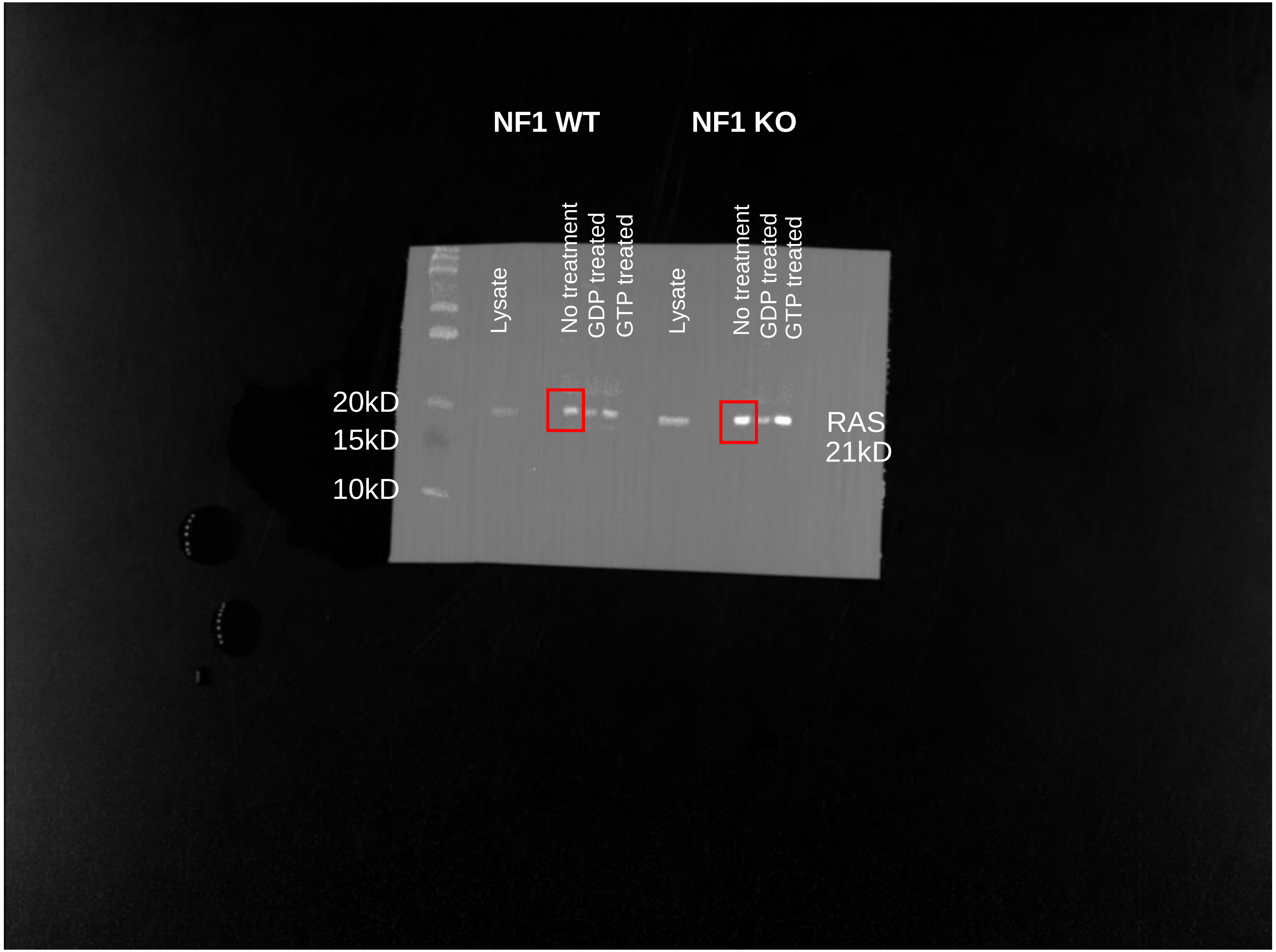

Supplementary Figure 2B.

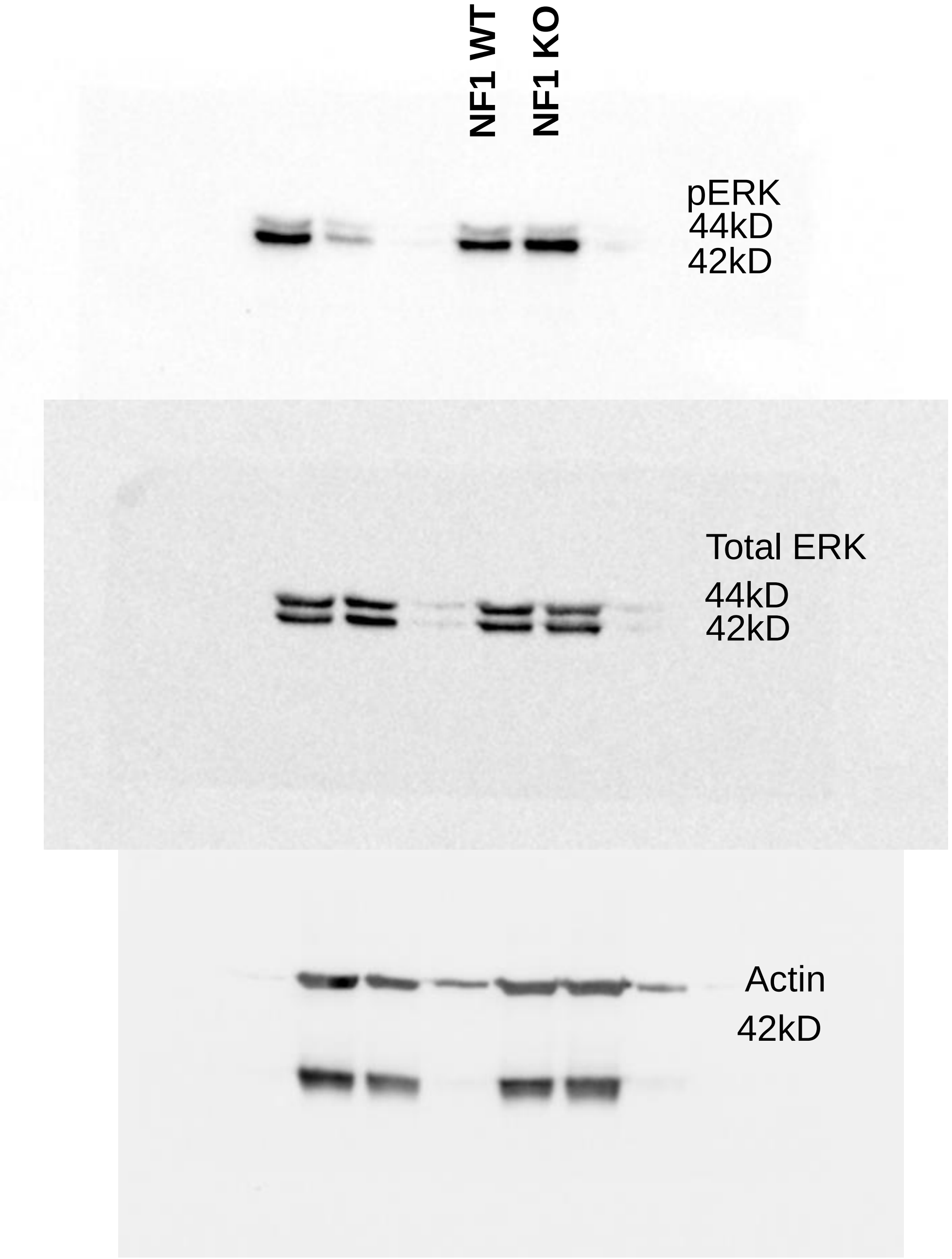

Supplementary Figure 3A.

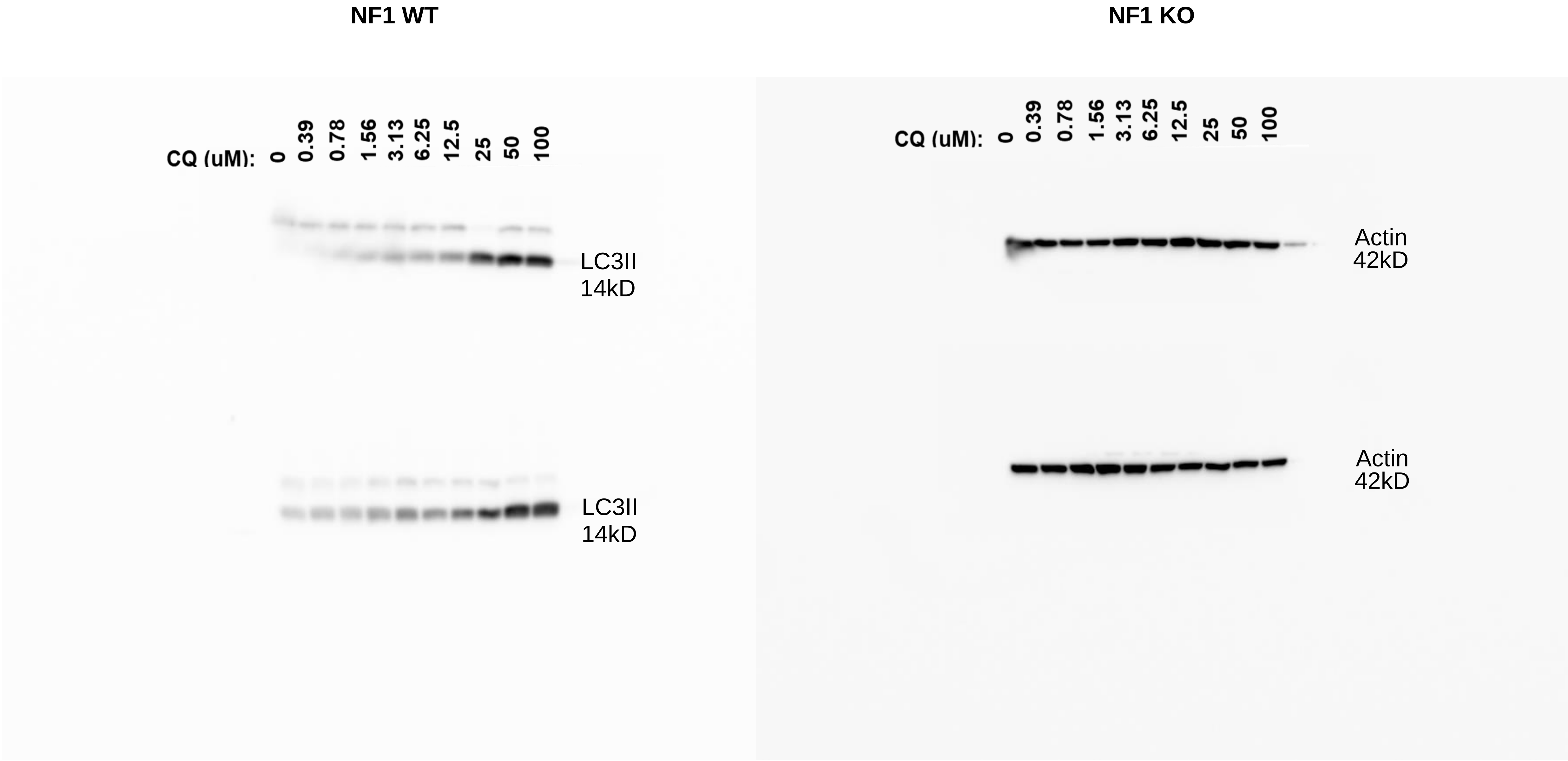

Supplementary Figure 3B.

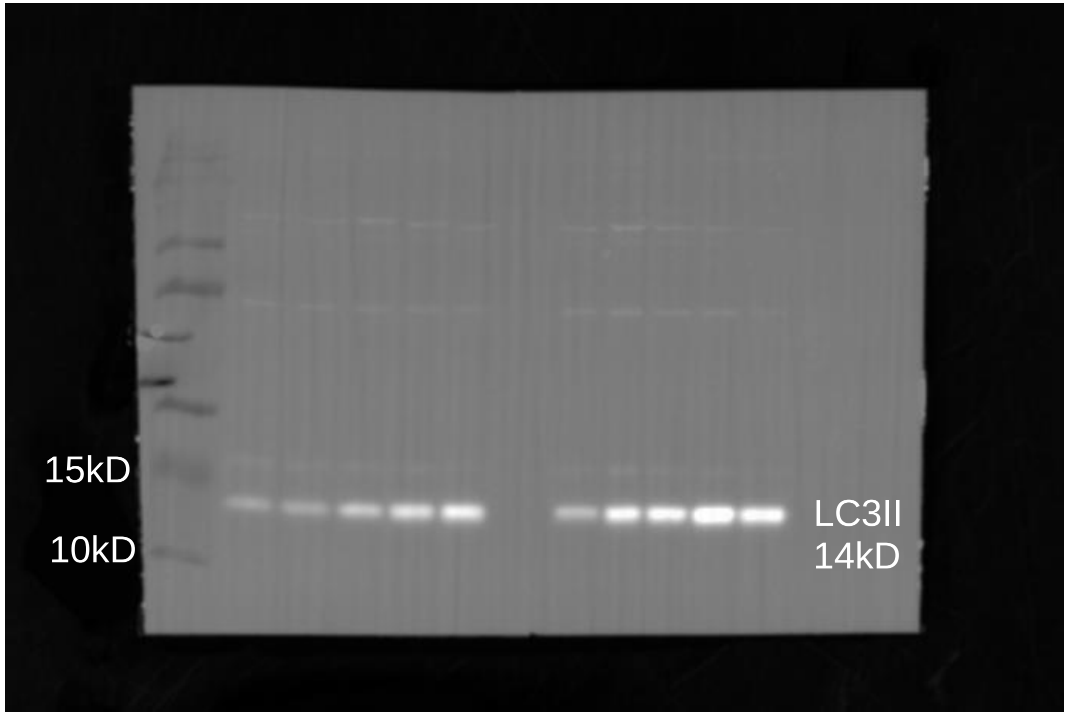


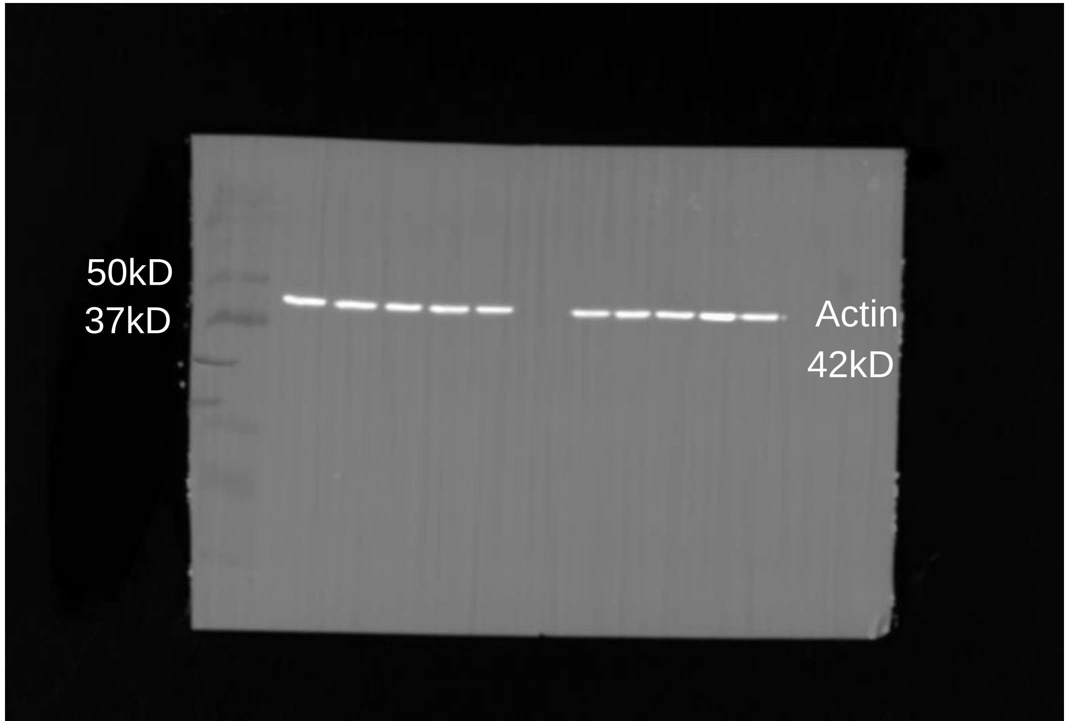

Supplementary Figure 3C.

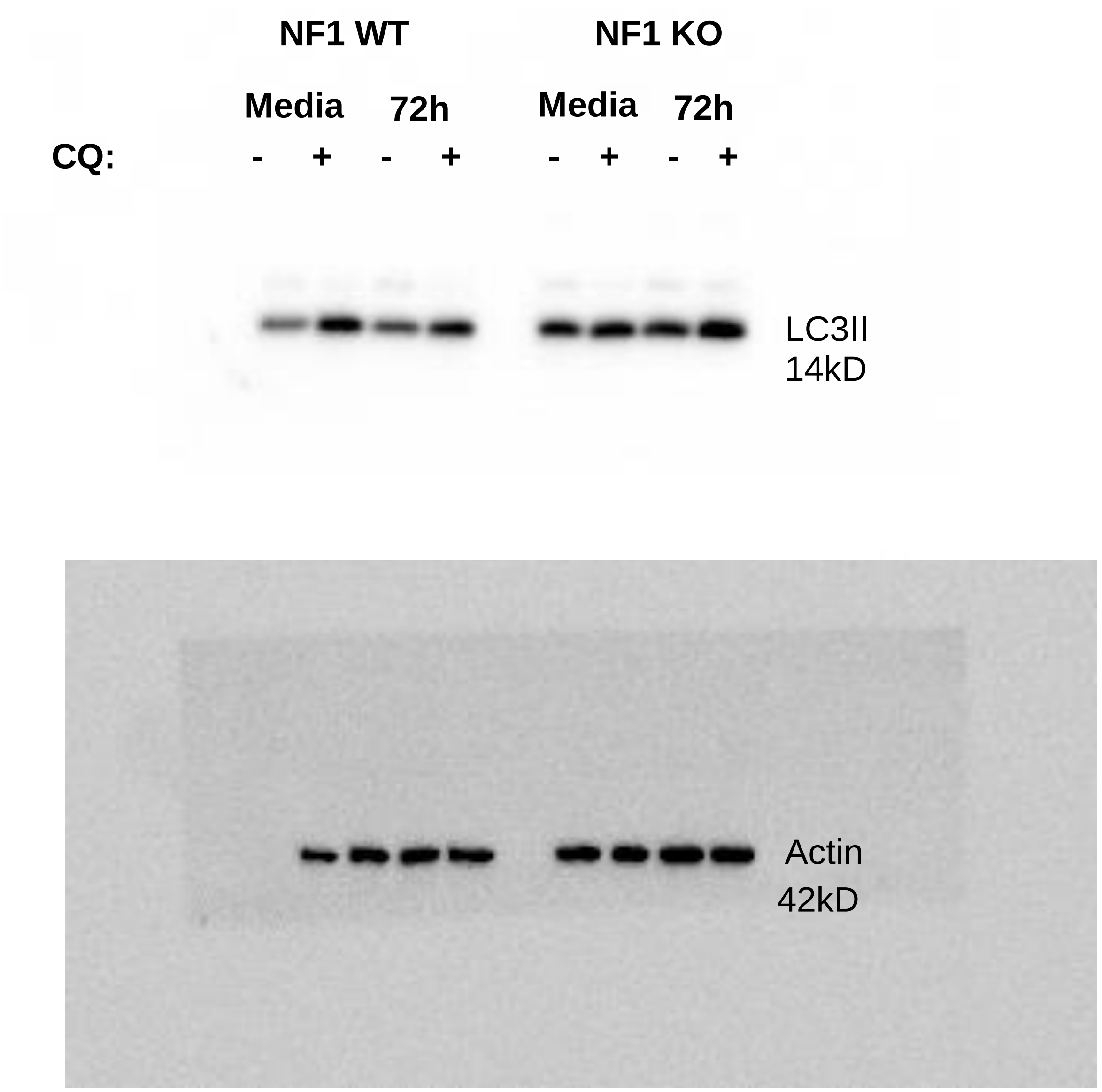
